# Accelerating the Validation of Endogenous On-Target Engagement and *In-cellulo* Kinetic Assessment for Covalent Inhibitors of KRAS^G12C^ in Early Drug Discovery

**DOI:** 10.1101/2022.02.17.480880

**Authors:** Vasudev Kantae, Radoslaw Polanski, Hilary J. Lewis, Derek Barratt, Bharath Srinivasan

## Abstract

Covalent inhibition is a valuable modality in drug-discovery due to its potential ability in decoupling pharmacokinetics from pharmacodynamics by prolonging the residence time of the drug on the target of interest. This increase in target occupancy is limited only by the rate of target turnover. However, a limitation in such studies is to translate the *in-vitro* inhibition assessment to the appropriate *in-cellulo* target engagement parameter by covalent probes. Estimation of such parameters is often impeded by the low-throughput nature of current label-free approaches. In this study, an ultra-performance liquid chromatography-multiple reaction monitoring (UPLC-MRM) mass spectrometry platform was utilised to develop a targeted proteomics workflow that can evaluate cellular on-target engagement of covalent molecules in an increased throughput manner. This workflow enabled a throughput increase of 5-10 fold when compared to traditional nanoLC-based proteomics studies. To demonstrate the applicability of the method, KRAS^G12C^ was used as a model system to interrogate the interaction of an irreversible covalent small-molecule, compound 25, both *in-vitro* and *in-cellulo*. Initial biochemical studies confirmed that the small-molecule forms an adduct with the targeted cysteine on the protein, as assessed at the level of both intact protein and on the target peptide. *In-cellulo* studies were carried out to quantify target engagement and selectivity assessment in heterozygous NCI-H358 cell line with both WT type and KRAS^G12C^ alleles. The workflow enabled evaluation of *in-cellulo* target engagement kinetics providing mechanistic insights into the irreversible mode of inhibition. In summary, the method has the potential for target agnostic application in the assessment of on-target engagement of covalent probes compatible with the high-throughput requirements of early drug discovery.

## Introduction

The resurgence of extensive exploration in small-molecules with covalent modality of action has spurred efforts at institution of methods for better characterization of irreversible inhibition^1^. Covalent inhibitors can prove highly advantageous in the open system that the human body represents. By optimizing for binding and subsequent reactivity-driven formation of the covalent adduct on the target of interest, the modality has the potential to uncouple pharmacokinetic clearance of a drug from its pharmacodynamic target engagement. Additionally, if resynthesis rate of the protein doesn’t compensate for target inactivation as a function of adduct formation, the desired effect will be long lasting. However, characterizing non-equilibrium irreversible modality of inhibition *in-vitro* and *in-cellulo* is challenging. Compounding this further, the persistent perception of off-target reactivity of covalent probes as a potential risk factor had muted the enthusiasm for exploring such drugs in the past. However, this perception is dissipating with demonstration of specificity and potency by several covalent inhibitors. A few prominent examples of traditional serendipitous covalent inhibitors include aspirin and other non-steroid anti-inflammatory drugs (NSAIDs) that inhibit the activity of cyclooxygenase (to treat inflammation and pain), sulphenamide modification of omeprazole for inhibiting H^+^/K^+^ ATP pumps (to treat gastroesophageal reflux and peptic ulcers), penicillin as an antibiotic that interferes with bacterial peptidoglycan synthesis and so forth. Of late, targeted electrophilic acrylamide warhead containing drugs (ibrutinib and afatinib) have been shown to target nucleophilic cysteine residues in BTK and EGFR,. Another prominent example of acrylamide warhead containing small molecule is Osimertinib (Tagrisso^®^, AZD9291), a covalent irreversible inhibitor of EGFR with preferential inhibition of sensitizing mutations and T790M resistance mutations (double mutants)^2^ over the wild-type receptor. More recently, FDA approved Sotorasib (Lumakras™) as an inhibitor of *KRAS^G12C^*, for patients with metastatic non-small cell lung cancer (NSCLC)^3^.

*In-vitro* characterization of irreversible inhibition has mainly relied on profiling^4^ and estimating the second-order parameter *k_inact_*/K_I_^5,6^ while *in-cellulo* efforts have been focussed on isolating and characterizing the adducts by approaches that use enrichment followed by nanoLC chromatography and high resolution mass-spectrometry. However, these efforts suffer from the disadvantage of being (semi)-quantitative, time-consuming and limited availability and hence, not compatible with early drug-discovery that relies on iterations of design and testing for optimization. An important challenge in early covalent probe discovery is the translation of biochemical parameters (*K_i_* or *k_inact_*/*K_I_*) into quantifiable cellular target engagement metrics. Medium to high-throughput assays are desirable to validate covalent biochemical hits for endogenous on-target engagement. These assays should be capable of (A) confirming on-target engagement with the desired target of interest, (B) confirming if compounds engages with the right nucleophile (e.g. cysteine) on the target and (C) enabling the ranking of compounds in cellular models. In addition, screening biochemical hits in a dose- and time-dependent manner across various cellular models is essential to determine the kinetics of on-target engagement. This will provide insights about the extent of target engagement that is required to elicit the desired biological outcome and for establishing a mechanistic link between efficacy and target inhibition.

As referred above, label-free methods employing nanoLC coupled to mass spectrometry (MS) are heavily employed to confirm target engagement. However these platform employ either a high-resolution mass spectrometer (typically Orbitraps and QTOF platforms), which are not easily accessible and are expensive, or are low-throughput approaches that employ nanoLC with typical run times of 30-90 min per sample^7–10^. Here we demonstrate the application of an Ultra Performance Liquid Chromatography (UPLC) and triple quadrupole (QQQ) mass spectrometer to develop a highly sensitive multiple reaction monitoring (MRM)-based targeted proteomics workflow to quantify the on-target engagement of covalent probes in cellular models. Traditionally, UPLC-QQQ has been widely used in targeted proteomics for biomarker screening applications owing to its advantages in demonstrating the sensitivity, specificity, reproducibility and, importantly, the throughput when compared with nanoLC-QTOF/Orbitrap based targeted proteomics^11,12^. Additionally, MRM based methods allow the absolute quantification of target proteins based on the selected proteotypic peptide(s)^13^. Larger diameter UPLC columns allow the injection of high sample volumes and thus, when combined with latest high sensitivity QQQ MS, these platforms allow accurate detection and quantification of medium-to high-abundance targets in cellular models.

As a proof-of-concept, the developed method was applied with KRAS^G12C^ as a model system to quantify the drug on-target engagement with a reported irreversible inhibitor of the protein^15^. Designing inhibitors against RAS has been a persistent challenge because of the high affinity of the protein for GTP and GDP (in the pM range), thus precluding the design of competitive inhibitors, and the lack of allosteric pockets with substantial geometric dimensions for effective inhibitor design. A number of investigators have demonstrated successfully that covalently targeting the mutated glycine to cysteine provides an tractable approach of targeting the variant, conferring the advantage of specificity and potency simultaneously^9,16,17^. However, several challenges remain that require novel solutions.

In summary, this article presents the implementation of a UPLC-MRM based methodology that enabled the quantification of *in-cellulo* target engagement kinetics with increased throughput for an irreversible inhibitor of KRAS^G12C^. We posit that this method has the potential to enable the mechanistic investigations of irreversible covalent inhibition in a target agnostic fashion. This methodological advance confers tangible advantages in making the SAR more reliable and for the translational potential of early drug-discovery efforts.

## Materials and methods

### Reagents and chemicals

All reagents and chemicals, unless mentioned otherwise, were procured from Sigma-Aldrich Co., (St. Louis, MO, USA), Amresco (Solon, OH, USA), or Fisher Scientific (Waltham, MA, USA). The experimental KRAS^G12C^ covalent inhibitor compound 25 was synthesised as described in (Kettle, Bagal et al. 2020). Cells were grown in RPMI1640 supplemented with 2 mM glutamine (Merck, Darmstadt, Germany) and 10 % Fetal Bovine Serum (ThermoFisher Scientific, Waltham, MA, USA). The cell-line NCI-H358 was obtained from the internal repository at AstraZeneca PLC and authenticated using Short Tandem Repeat (STR) fingerprint analysis and were confirmed mycoplasma free. Recombinantly expressed and purified KRAS^G12C^ protein was provided by the Protein Science group, AstraZeneca PLC. Anti-RAS antibody was purchased from Merck Millipore. Pierce™ recombinant protein A/G and clear round-bottom immuno nonsterile 96-well plates were obtained from ThermoFisher Scientific. Stable isotopically labelled (SIL) human KRAS^G12C^ (isoform 2B) protein was purchased from Promise Advanced Proteomics. Sequence grade modified Trypsin (Porcine) was ordered from Promega UK Ltd, Southampton Hampshire, UK.

### Bioinformatics and structure analysis

The sequence of WT KRAS, HRAS and NRAS and its variants (including G12C) were obtained from the repository at NCBI. The non-redundant database at NCBI was used to search for homologs of KRAS using the algorithm blastp. t-coffee^18^ was used to generate multiple sequence alignment profiles and the figure was generated using ESPRIPT^19^. The sdf file of the smallmolecule (compound 25) was used for analysis and the structure image was generated using ChemDraw version 20.0. Structures of protein solved using x-ray crystallography, were downloaded from PDB (accession codes: 6t5b, 6t5u and 6t5v). Molecular visualization and structure analysis were performed using spdbv^20,21^ and pymol (pymol.sourceforge.net)^22^.

### Sample preparation

#### A. In vitro biochemical reaction

Hexahistidine-tagged KRAS^G12C^ and KRAS^WT^ proteins (isoform 2B, residues 1–169, based on the construct used for PDB 3GFT) were recombinantly expressed in *Escherichia coli*, purified and complexed with GDP at Protein Sciences Department within AstraZeneca PLC.

#### B. Covalent Engagement Assays

KRAS^G12C^ protein at concentration of 200 nM was incubated at RT with compound 25 at three different concentrations (0.1, 1 and 10 μM) in buffer containing 20 mM bicine pH 7, 100 mM NaCl for 4 hrs or at different time points (0, 60, 120, 180 and 240 minutes, respectively). The reaction was quenched with reducing buffer (20 mM DTT in H_2_O). The samples were then denatured by incubating at 60 °C for 30 min, followed by alkylation with iodoacetamide in dark for 20 min. Samples were digested with sequence grade trypsin (1:25 ratio of trypsin to sample) for 16 hrs at 37 °C, followed by drying the samples in speedvac and reconstituting in 0.1 % TFA. 10 μl of samples were injected for LC-MRM analysis, as described below.

#### C. Intact mass analysis

KRAS^G12C^ protein at concentration of 1 μM was incubated at 37° C with covalent inhibitor at 0, 0.5,1 and 10 μM in buffer containing 50 mM ammonium bicarbonate pH 7.5, 50 mM NaCl for 2.5 hrs and the reaction was quenched with 1% (v/v) formic acid, followed by analysis in RapidFire-QTOF (Agilent Technologies, Palo Alto, CA, USA), as described below.

### Cellular KRAS^G12C^ target engagement samples

NCI-H358 cells (2×10^6^) were seeded into the wells of six-well plates (Corning) in 2 mL of culture media, cultured overnight and treated with the covalent compound at three different concentrations (0.1, 0.3 and 1 μM) and harvested at five time points (0, 1, 2, 3 and 4 hrs) by first washing twice with ice-cold PBS and adding 100 μl/well of the lysis buffer [20 mM Tris HCl, pH 7.4, 137 mM NaCl, 10 % (v/v) glycerol, 1 % (v/v) Triton X-100, 0.5 % (w/v) sodium deoxycholate, 0.1 % (w/v) SDS, 2 mM EDTA, tablet of protease and PhosStop inhibitor cocktail (Sigma-Aldrich) and Bezonase inhibitor]. Lysates were transferred to 1.5 ml Eppendorf tubes, centrifuged and the supernatant was collected to fresh tubes. Total protein concentration in each sample was measured using BCA assay and normalized to 1 mg/ml concentration.

An immunoprecipitation based protocol based on previous report was used to enrich KRAS^G12C 15^. Briefly, immuno nonsterile 96-well plates were coated with protein A/G and then Anti-RAS antibody (**Figure 1A**). 25 μg of cell lysate was added and spiked with SIL KRAS^G12C^ protein (Lys-^13^C_6_,^15^N_2_ and Arg-^13^C_6_,^15^N_4_) (4 pmole per 100 μg of total protein), as an internal standard. Following immunoprecipitation, samples were sequentially washed with RIPA buffer (Sigma), phosphate buffered saline and water. Samples were reduced with DTT, alkylated with iodoacetamide (IAM) and trypsin digestion was carried out 18h at 37 °C. Samples were dried by centrifugal evaporation and reconstituted in 0.1 % TFA for UPLC-MRM MS analysis.

**Figure1:**
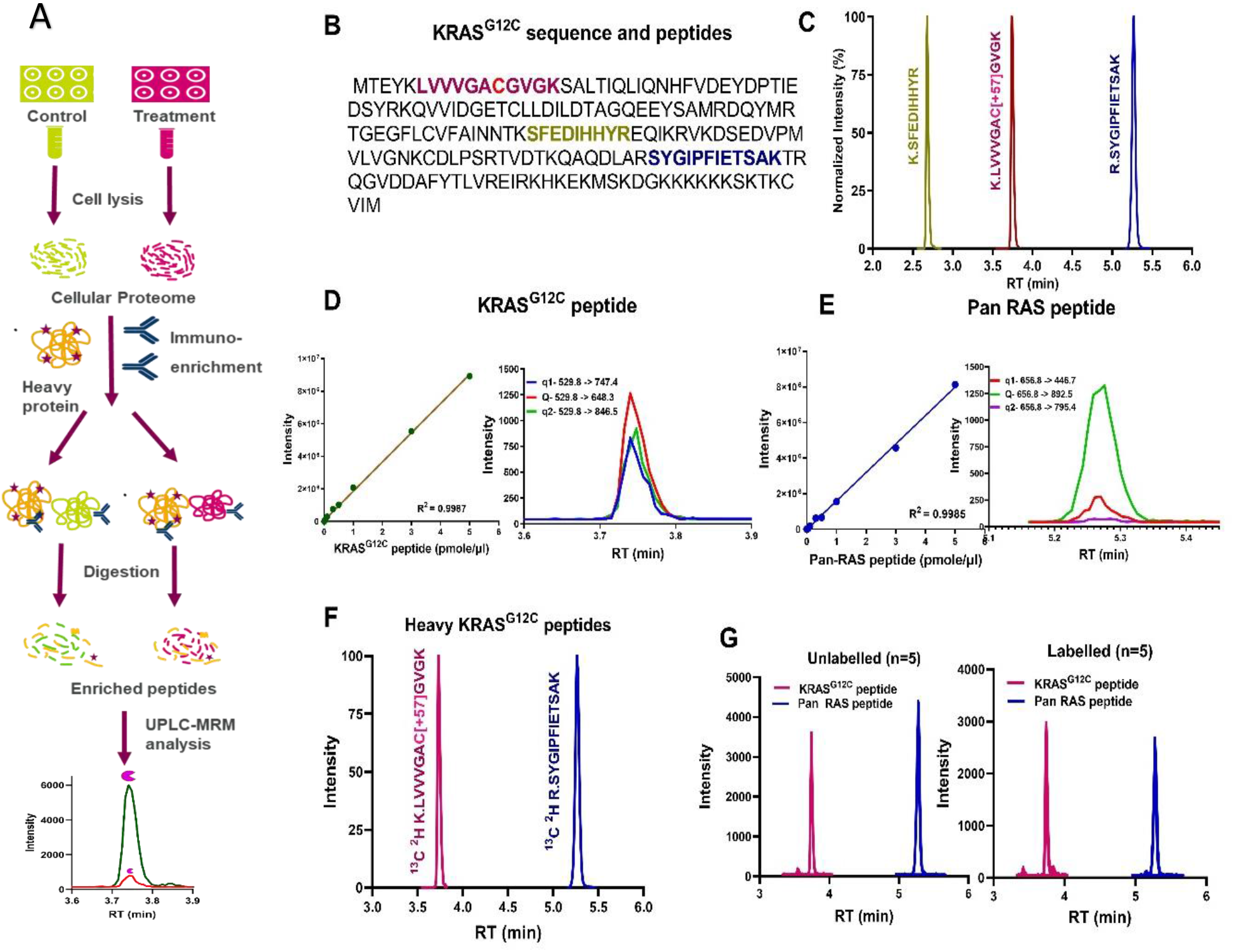
Analytical method development and validation parameters. **(A)** immuno-enrichment based sample preparation flowchart. **(B)** KRAS^G12C^ sequence and the coloured are the peptides selected for method development. **(C)** Chromatogram displaying the separation of the selected three peptides using recombinant protein digest. **(D&E)** Calibration curves of G12C and Pan RAS peptides showing the linear range up to 4 orders of magnitude. Chromatograms showcasing the limits of detection (LODs) levels of G12C peptide and Pan RAS peptides, 300amole and 15fmole on column respectively. The LODs are calculated by S/N>3 and by visual inspection. Three different MRM transitions were selected for each peptide, of which the intense one is used as quantifier (Q) and other two as qualifiers (q). Red (529.8>648.3) and green (656.8>892.5) coloured transitions were used as Q for G12C and Pan RAS peptides respectively. **(F)** Chromatograms of SIL G12C and Pan RAS peptides. The retention time of these are similar to the unlabelled from of their peptides. **(G)** Chromograms showcasing the method precision for both G12C (magenta) and Pan RAS endogenous peptides (blue) and their labelled variants spiked before sample processing. The method precision is displayed in %RSD (relative standard deviation) and for both peptides the RSD is <5%.

### UHPLC-MRM MS analysis

The targeted analysis was performed on a standard-flow UPLC-MS platform using an Agilent 1290 Infinity UPLC system and an iFunnel Agilent 6495c QQQ mass spectrometer (Agilent Technologies, Palo Alto, CA, USA), fitted with a standard-flow ESI (Jet Stream) source. The UPLC system consisted of a reverse phase (RP) chromatographic column (50 × 2.1 mm. i.d., Agilent Zorbax Eclipse Plus C-18 Rapid Resolution HD, 1.8 μm particle), maintained at a temperature of 45 °C. Separation of peptides was achieved at flow rate of 0.4 mL/min using a multi-step gradient with mobile phase A (0.1 % FA v/v) and mobile phase B (acetonitrile 0.1 % FA v/v) as follows (time, %B): 0 min, 3 %; 0.5 min, 3 %; 5 min, 25 %; 6 min, 40 %; 6.5 min, 95 %; 7.2 min, 3 %, and a post-column cleaning for 1.3 min.

The Agilent QQQ mass spectrometer was operated in the positive ion mode. The MRM acquisition parameters were 3,500 V capillary voltage, 300 V nozzle voltage, 11 L/min sheath gas flow (UHP nitrogen) at a temperature of 250 °C, 15 L/min drying gas flow at a temperature of 150 °C, 30 PSI nebulizer gas flow, iFunnel voltages (150 V and 50 V at high and low pressure) and unit resolution [0.7 Da full width at half maximum (FWHM)) in the first quadrupole (Q1) and the third quadrupole (Q3). The default fragmentor voltage of 380 V and a cell accelerator potential of 4 V was used for all MRM ion pairs, and the dynamic MRM option was used for data acquisition. **Table 1** shows the MRM transitions selected for each peptide.

**Table 1.** MRM transition of light and heavy peptides

| Species | Sequence | Q1 (m/z) | Q3 (m/z) |
| --- | --- | --- | --- |
| KRAS<sup>G12C</sup> | LVVVGAC[+57.0]GVGK | 529.805 | 648.3 |
|  |  |  | 747.4 |
|  |  |  | 846.5 |
| KRAS<sup>G12C</sup> heavy | LVVVGAC[+57.0]GVGK [+8] | 533.8 | 656.3 |
|  |  |  | 755.4 |
|  |  |  | 854.5 |
| RAS WT | LVVVGAGGVGK | 478.3 | 644.6 |
|  |  |  | 545.3 |
|  |  |  | 743.4 |
| Pan RAS | SYGIPFIETSAK | 656.8 | 446.7 |
|  |  |  | 795.4 |
|  |  |  | 656.8 |
| Pan RAS heavy | SYGIPFIETSAK [+8] | 660.8 | 450.7 |
|  |  |  | 803.7 |
|  |  |  | 900.5 |

### RapidFire-QTOF analysis of intact protein

An Agilent RapidFire 365 model was used to process and inject samples. Briefly, all solvents are delivered using three Agilent 1200 Quaternary pumps. A single C4 Type A (Agilent, G9203) SPE cartridge was used for the sample clean-up. 10 μL samples were aspirated for 600 ms from each well of a 384-well plate. The sample load/wash time was 5000 ms at a flow rate of 1.25 mL/min (95/5 H_2_O/ACN, 0.1 % v/v FA) and elution time was 7000 ms 70/30 ACN/H_2_O, 0.1 % v/v FA and finally with a re-equilibration time of 600 ms at a flow rate of 1.25 mL/min (H_2_O, 0.1 % v/v FA). The overall analysis time per sample was 18 s/well. Methanol and water were used as organic and aqueous washes to clean the needle post-injection.

Agilent iFunnel 6550 QTOF (Agilent Technologies, Palo Alto, CA, USA) was used to acquire the intact protein mass in positive ion mode, operated in electrospray ionization using dual jet stream. The instrument parameters were as follows: gas temperature 350 °C, drying gas 5 L/min, nebulizer 60 PSI, sheath gas 350 °C, sheath gas flow 11 L/min, capillary 3.5 kV, nozzle 0 kV, fragmentor 200 V, funnel voltage at 150 and 60 V at high and low pressure. Data were acquired at a rate of 5 spectra/s. The mass range was calibrated using the Agilent positive ion tune mix, over the *m/z* range 300–3200. The instrument control software was Agilent MassHunter Workstation Data Acquisition.

### UPLC-QTOF peptide mapping analysis

The MS/MS analysis of the peptide for sequence confirmation was performed on a UPLC coupled to Agilent iFunnel 6550 QTOF mass spectrometer. The UPLC parameters and the QTOF source parameters were as described above. Data were acquired in auto MS/MS mode, with mass range of 110-1700 m/z at acquisition rate of 8 spectra/sec and MS/MS with mass range of 50 to 1700 m/z at 3 spectra/sec. Differential collision energy was applied for 2, 3 and >3 charge states.

### Data analysis and statistical validation

All MRM data were processed with Agilent’s MassHunter Quantitative and Qualitative Analysis software. Mass hunter quantitative software was used to build calibration curves and quantitate the absolute concentration of peptides in the samples. Response ratios (area of unlabelled peptide/area of SIL peptide) were used for each peptide to determine these concentrations. The KRAS^G12C^ peptide area for each sample was normalised to the pan RAS peptide signal. Target engagement was calculated as a percentage relative to the vehicle control (Scheme I). Agilent Bio-confirm suit was used to deconvolute the raw spectrum to determine the accurate *m/z* of the protein and peptides. Processed data was statically analysed in GraphPad Prism (v9).

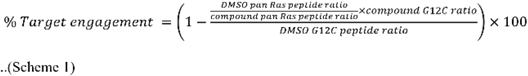

### Kinetic analysis for in-cellulo target engagement data

A covalent inhibitor binds to the enzyme and forms an irreversible bond with the enzyme of interest (Scheme I). The sequence of events involve binding of the inhibitor and subsequent inactivation. This is evident as a time-dependent reduction in the rate of the reaction from initial velocity (*v_i_*) to steady-state velocity (*v_ss_*). The *v_ss_* is usually zero for covalent irreversible inhibition.

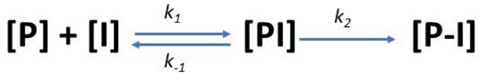

Where, P and I are free protein and inhibitor respectively, PI is the non-covalent complex between protein and inhibitor and P-I is the covalent complex between the protein and the inhibitor. Since we are looking at loss of peptide from apo-protein as an indirect report on adduct formation in this study, the rate constant *k_2_* is the maximal rate of modification (*k_mod_*). *K_I_*, inhibitor concentration at half maximal modification rate when all the inhibitor is complexed with enzyme, is defined as (*k*_−*1* +_ *k_2_*) *k_1_*. In the current study, *k_mod_* can be directly equated with *k_inact_* since only a single alkylation event was observed and it is highly likely that modification is directly proportional with inhibition. However, this could not be assessed in the absence of a biochemical assay. Complete progress curves were generated in NCI-H358 cellular model at several different concentration of compound 25 and were fitted to equation 1. Note that *V_ss_* is zero indicating maximal inhibition as is expected for covalent inhibitors.

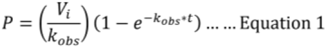

The *k_obs_* estimated from this fit was plotted as a function of inhibitor concentration and the resultant data points were fit to the equation 2 for a line.

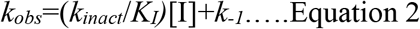

The slope of the line estimates the second order parameter *k_inact_*/*K_I_* with units of concentration inverse time inverse.

It would have to be reiterated here that the *k_obs_* versus [I] replots can either be linear or hyperbolic depending on the relative magnitude of the *k_−1_* and *k_2_* terms. Hyperbolic replots enable the resolution of *K_I_* and *k_inact_* terms while linear replots enable the estimate of their ratio (*k_inact_*/*K_I_*). The range of *k_−1_* and *k_2_* values that results in either a hyperbolic or linear secondary replots has been discussed previously^23^).

## Results

### Optimization of UPLC-MRM analysis

A highly sensitive and robust method is a pre-requisite to accurately quantify the target engagement in cellular samples. This section explains the method development, which was initially optimized to assess biochemical target engagement and subsequently applied to cellular samples (**Figure 1A)**. Recombinant KRAS^G12C^ protein digest was used to optimize the UPLC and mass-spectrometer parameters. Agilent Skyline automation tool^24^ was used to create methods for analysis of peptides from recombinant KRAS^G12C^ protein (**Figure 1B**). The sequence was imported into Skyline for *in-silico* tryptic digestion and the methods were directly exported to the instrument for acquisition. The results generated were imported back into Skyline for manual inspection and this process was repeated iteratively to arrive at the final method. Three proteotypic peptides, including the peptide with the target cysteine, were selected (**Figure 1B and 1C**). LVVVGA**C**GVGK is the unique proteotypic peptide containing cysteine at codon 12 and is specific to the KRAS^G12C^ isoform. SYGIPFIETSAK is used as a Pan RAS peptide (conserved in other RAS proteins e.g. NRAS, HRAS). The three most sensitive MRM transitions for each peptide were screened for matrix interference. This is essential due to the complexity of the proteome and the low resolution mass spectrometer used for the targeted analysis. The high intensity transition is used as the quantifier (Q) ion and other two transitions were used as qualifier ions (q), respectively. The quantifier to qualifier ion ratio from earlier experiments was used to monitor the increase in specificity of the peptide in endogenous samples (**Figure 1D and Figure 1E**). The UHPLC gradient was optimized to achieve chromatographic separation between peptides and to enable high-throughput. System parameters were further optimized to ensure no carryover between injections and to maintain analytical robustness of retention times. SIL KRAS^G12C^ protein digest was subject to the same process to determine optimal MRM and LC parameters. The transitions and retention time (RT) were similar to unlabelled peptides **(Figure 1F)**. Additionally, no traces of unlabelled peptides were found in the labelled peptides samples.

The developed method had a detection limit (LOD) of 300 attomole on column for the KRAS^G12C^ peptide and 15 femtomole on column for Pan RAS peptide and provided a linear range up to 4 orders of magnitude (**Figure 1D and Figure 1E**). The method was robust resulting in high precision (Relative Standard deviation RSD < 5 %) across multiple runs. **Figure 1G** depicts the precision of endogenous KRAS^G12C^ and Pan RAS peptide, along with spiked SIL peptide (n-=5 biological replicates).

### Biochemical interrogation of target-small molecule interaction

RAS is a superfamily of proteins with highly conserved N-terminus as seen in the multiple sequence alignment (**Figure S1**). In a previous report, it has been shown that irreversible small-molecules target an allosteric pocket on the GDP-bound KRAS^G12C^, thus, locking it in an inactive state^15^. A series of piperazine-quinazoline motif containing small-molecules provided the best inhibition with enhanced selectivity and potency against the mutant isoform of KRAS as shown in the structural superposition of the respective PDB structures (PDB IDs: 6t5b, 6t5u and 6t5v, respectively) (**Figure 2A**). In this study, compound 25 (**Figure 2B**) was selected as the most potent of the series, to carry out our mass-spectrometry based studies on target engagement *in-vitro* and *in-cellulo*.

**Figure 2.**
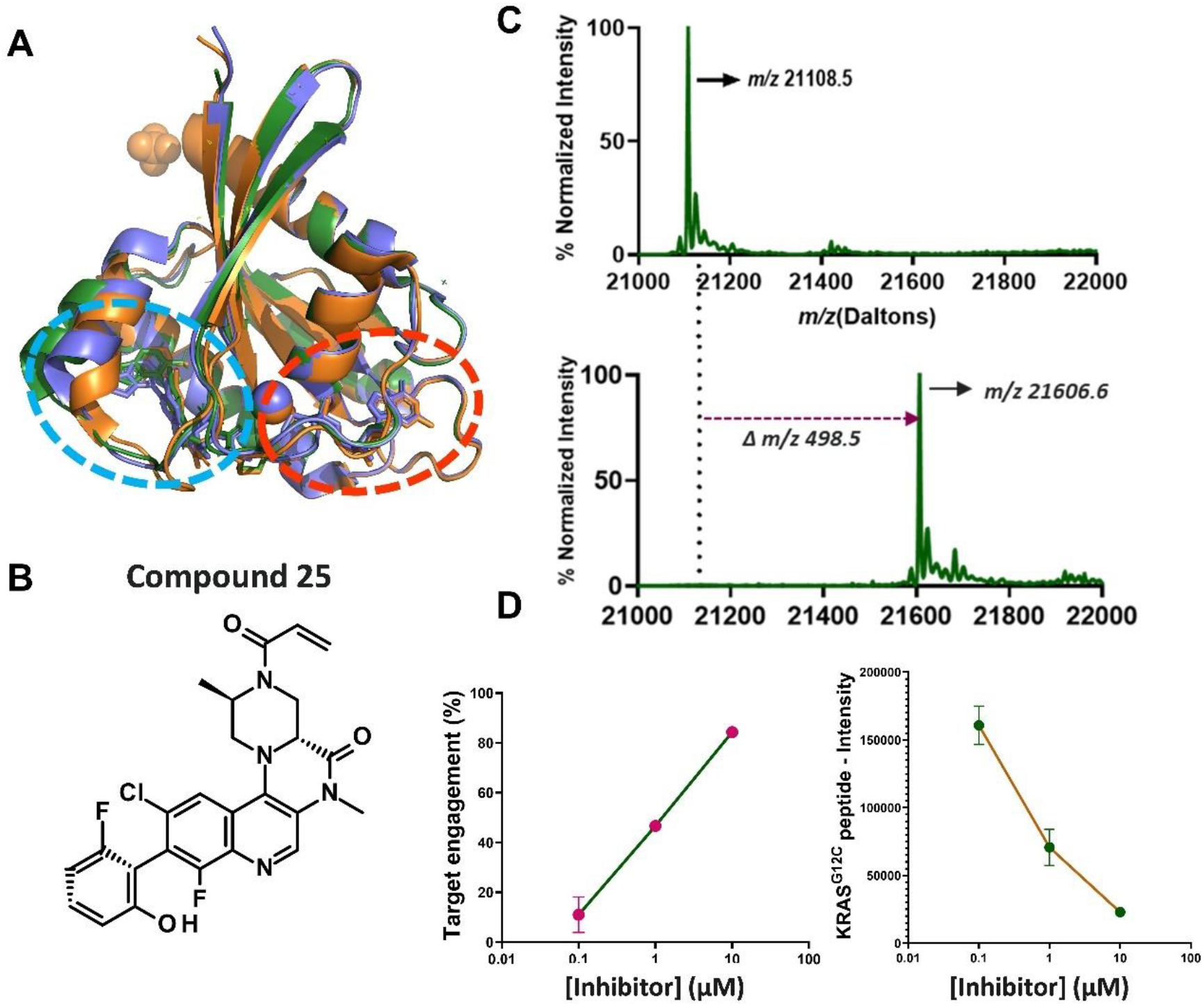
Biochemical interrogation of adduct formation **(A)** Structural superposition of co-crystallized structures reported ^15^ showing the position of the allosteric inhibitor binding pocket (cyan dotted line) and the GTP binding pocket (red dotted line). The PDB id of the structures are 6T5V(Green), 6T5B (Orange) and 6T5U (Purple). **(B)** Structure of the small-molecule inhibitor (compound 25)^15^ employed in this study for assessment of target engagement **(C)** Target engagement interrogated at the intact protein level. The zero-charge Maximum Entropy deconvoluted MS spectra of KRAS^G12C^ protein at 1μM. Upon incubation for 2.5h with the small-molecule at 1 μM concentration, the *m/z* for the target protein shifts by 498.9 Daltons corresponding to the molecular mass. **(D)** Biochemical target engagement assessed for the unique KRAS^G12C^ peptide as a function of increasing concentration of inhibitor incubated for 4h. The fraction target engagement was computed by normalizing for the intensity of the pan RAS peptide as specified in materials and methods section.

As a first step, *in-vitro* characterization of adduct formation was undertaken both with intact protein and peptide based approaches. Intact protein analysis was carried out that confirmed that the average *m/z* of the KRAS^G12C^ protein (*m/z* of 21108.2 Da) was similar to the theoretically computed value (**Figure 2C**) (the multiply charged ions in the mass spectrum are shown as **Figure S2**). Further, incubation with compound 25 (average *m/z* 499.1 Da) resulted in the formation of single adduct at *m/z* of 21607.1Da (**Figure 2C**). The *m/z* of compound 25, forming the irreversible adduct with the protein, was confirmed by injecting the compound alone **(Figure S3).**

Next, tryptically digested recombinant KRAS^G12C^ samples were injected to confirm the sequence of the selected peptides by performing MS/MS analysis **(Figure S4A**). The same analysis was repeated after preincubation of the protein with the compound 25. Results indicate that the compound forms an adduct with the target protein and depicts a single binding site. To further confirm if the adduct formed by compound 25 occurs with the cysteine at position 12, incubated biochemical samples were peptide mapped by MS/MS analysis (**Figure S4B**).

After confirming that the irreversible small-molecule inhibitor was forming an adduct with Cys12, target engagement levels were quantified in a dose-dependent manner using the developed UPLC-MRM QQQ method. A dose-dependent increase of target engagement was observed when the compound was incubated for 4 hrs at room temperature (**Figure 2D**). Similarly, a dose-dependent decrease of the KRAS^G12C^ peptide was quantified demonstrating that an increase in adduct formation correlates with a near simultaneous decrease in the non-modified peptide (**Figure 2D**). This *in-vitro* characterization of the adduct formation formed the basis for subsequent *in-cellulo* target engagement kinetics.

### Interrogation of in-cellulo target engagement in NCI-H358 cellular model

After the *in-vitro* validation, the method was applied to quantify target engagement of endogenous KRAS^G12C^ and Pan RAS peptides by the compound 25 in cellular model. NCI-H358 cells were selected for these studies since this NSCLC line harbours a heterozygous missense KRAS mutation at codon 12 constitutively activating the EGFR-KRAS pathway. When NCI-H358 cells were treated with the covalent compound for 4 hrs, the target peptide levels depleted substantially. This was similar to *in-vitro* observations (see section 3.2). Quantification of the levels indicated 96 % target engagement (**Figure 3A**). SIL protein was spiked into cell lysates before sample processing to correct for losses and to determine absolute concentration. The unaltered levels of SIL protein further confirmed the endogenous specificity of the monitored peptides. To demonstrate that the observed target engagement is not an artifact of monitoring the KRAS^G12C^ peptide but is exclusively compound mediated effect, DMSO and compound treated samples were diluted in different ratios to enable a dose-dependent increase in compound concentration that translates into different levels of target engagement. **Figure 3B** shows an increase in target engagement with increase in inhibitor to DMSO ratio, which further demonstrates the analytical validation of the developed method in capturing different levels of target engagement in cellular samples.

**Figure 3.**
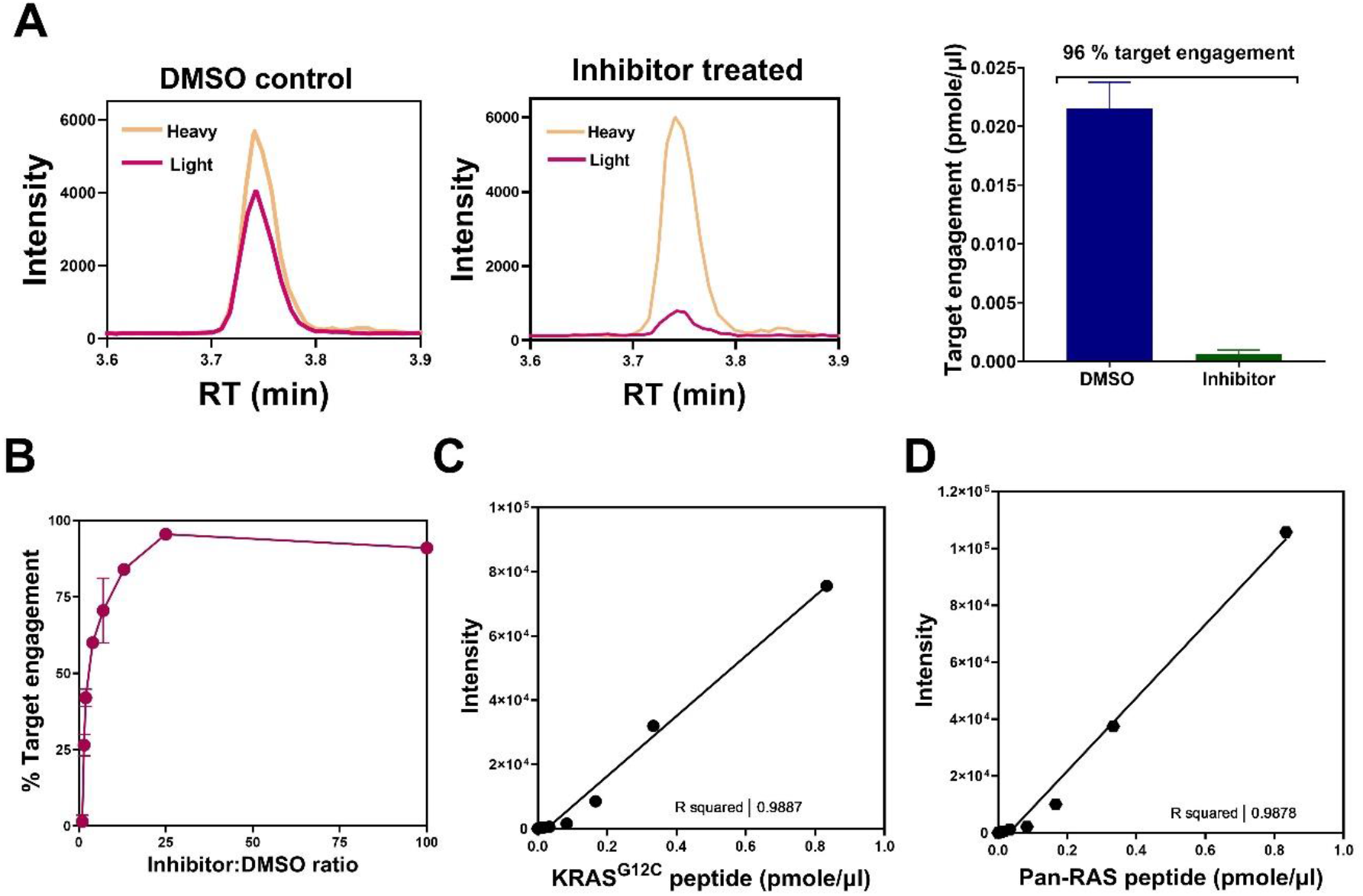
*In-cellulo* target engagement. **(A)** Chromatogram displaying the loss of G12C peptide (red) when NCI-H358 cell samples were treated with 10μM of compound 25 for 4h at 37°C. The SIL variant of G12C peptide (yellow) is unaffected. The absolute amount of target inhibition is shown in histogram. **(B)** Target engagement as a function of increase in inhibitor to DMSO ratios. DMSO and compound treated samples were diluted in different ratios to capture different levels of target engagement. (**C & D**) Endogenous calibration curves of G12C and Pan RAS peptides generated by spiking known amount of SIL protein to NCI-H258 cell lysates.

SIL protein at different starting concentrations was spiked prior to sample preparation to show that the method had linear response in presence of matrix. Both KRAS^G12C^ and Pan RAS peptide shows an endogenous linear response of up to 2-orders of magnitude (**Figures 3C** and **3D**).

### Kinetics of cellular target engagement

Often, in early drug discovery, the principal challenge is to translate and correlate the *in-vitro* kinetic observations with those obtained *in-cellulo* in disease relevant models^25^ **(Figure 4A**). The rate limiting step here is a method that can report back on *in-cellulo* target engagement without relying on biomarkers that can distort the kinetic readout. With the aim of bridging this gap using the label-free method developed above, a dose and time-dependent response was generated for the inhibition of KRAS^G12C^ by the compound 25 in NCI-H358 cells. Cells were treated with covalent inhibitors at three different doses (0.1 μM, 0.3 μM and 1μM) across time intervals spanning 0, 1, 2, 3 and 4 hrs to quantify target engagement. As is seen in **Figure 4B**, a clear increase of target engagement with increase in dose and time is evident from the area under the curve. The SIL peptides were not effected as a function of increasing dose (**Figure S5**). Maximal engagement was achieved between 3-4 hrs at a 1 μM dose of the inhibitor. In addition, given the heterozygous nature of the NCI-H358 cellline with both KRAS^G12C^ and WT alleles, we aimed to understand whether the method has capability to individually monitor and discriminate between the WT and the G12C peptide, thus helping to determine compound selectivity. As shown in **Figure 4B** (top panel), though a clear dose-dependent inhibition is observed for KRAS^G12C^ peptide when treated with compound 25, the compound treatment has no effect on the WT peptide even when treated at high doses for 4 hrs. In combination of the biochemical inferences, this provides strong evidence for specific targeting of the cysteine at the 12^th^ position of KRAS^G12C^ *in-cellulo* and discounts the possibility that non-specific reactivity is contributing to adduct formation. Further, this result paves the way for the interrogation of *in-cellulo* target engagement in a target agnostic way at high resolution and increased throughput.

**Figure 4.**
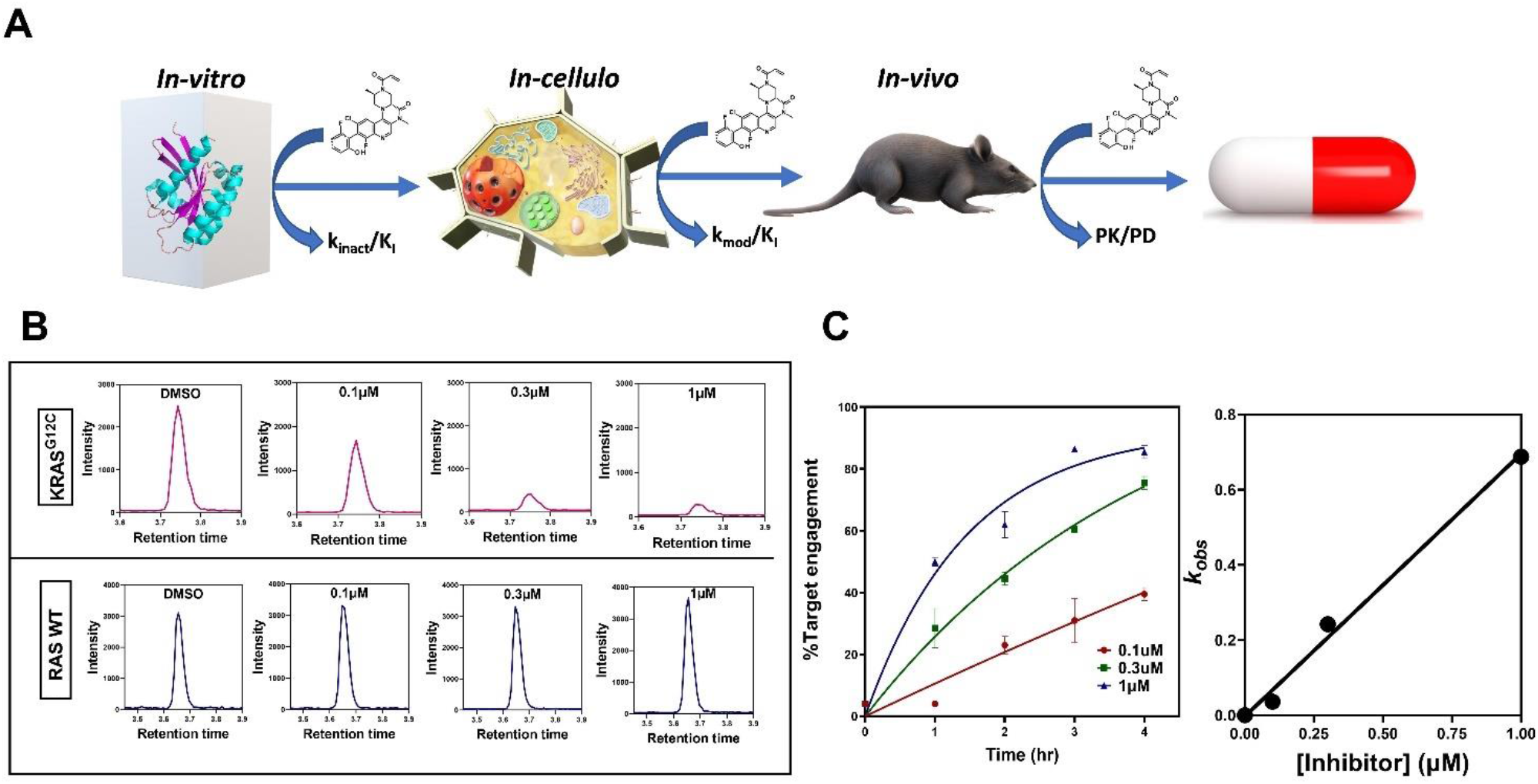
Kinetics of cellular target engagement **(A)** Schematic representation of the translatability of parameters estimated *in-vitro* to *in-cellulo* and, finally, to *in-vivo* models. **(B)** Concentration-dependence of *in-cellulo* target engagement for KRAS^G12C^ and demonstration of selectivity against the variant vis-à-vis the WT protein in a heterozygous NCI-H358 cell line **(C)** Progress curves generated at different concentrations of the inhibitor were fitted to models for irreversible inhibition to estimate the parameter *k_obs_*. Secondary replot of *k_obs_* versus inhibitor concentration yielded the second order parameter *k_inact_*/*K_I_*.

Data obtained from experiment shown in Figure 4A allowed us to estimate kinetic parameters. The data were replotted as progress curves across several doses of compound 25 for % target engagement (**Figure 4C**) and as the absolute concentration of KRAS^G12C^ peptide (**Figure S6**). Since the compound 25 is a covalent and irreversible inhibitor of KRAS^G12C^ targeting the nucleophilic cysteine, it was expected that the progress curves would be non-linear. This is because non-equilibrium slow-onset or irreversible inhibitors show time-dependence of inhibition apart from dose dependent behavior^26^. As evident from perusal of the progress curves in **Figure 4C**, the non-linearity of the endogenous target engagement curves, showing transition from an initial velocity (*v_i_*) to a steady state velocity (*v_ss_*) as a function of time-dependent inhibition, is clear. The rate of transition from *v_i_* to *v_ss_* was estimated by fitting to the equation 1 specified in materials and methods (**Figure 4C**). The resultant values of *k_obs_* was plotted versus the inhibitor concentration and the data points were fitted to an equation for line (equation 2) with the slope yielding us the value for *k_mod_*/*K_I_* **(Figure 4C)**. It would have to be noted that *k_mod_* is different than *k_inact_*. To establish parity between the two parameters, a correlation would have to be established between modification and inactivation. This might not be always possible, especially for proteins that do not have an enzymatic activity or an assay that will enable one to assess inhibition: see materials and methods for further details). A value of 248 ± 73 M^−1^ sec^−1^ (arithmetic mean ± standard deviation) was estimated for the *in-cellulo k_mod_*/*K_I_*. The parameter was measured with experiments performed on biological duplicates on two different occasions and two technical replicates each, respectively. This is the lower limit for the estimate of *k_mod_*/*K_I_*. This is because the value is estimated by plotting the total concentration of the inhibitor as a function of *k_obs_*. However, under cellular assay conditions, the total inhibitor concentration could be markedly different than the inhibitor concentration available to interact with the target of interest due to issues of uptake, macromolecular crowding, non-specific sequestration of the inhibitor, inhibitor degradation (if any) and establishment of equilibrium between the media and cell. To the best of our knowledge and belief, this is the first attempt at estimating the *k_mod_*/*K_I_* parameter for *in-cellulo* target engagement kinetics using mechanistic models and approaches optimized for estimation of biochemical *k_inact_*/*K_I_*.

## Discussion

Understanding target engagement intracellularly, especially during early stages, is an important criterion for driving the success of a drug discovery program (**Figure 5**). Inferential approaches such as target engagement *in-vitro* (assessment using biochemical assay or x-ray crystallographic structure solution) translating into improved therapeutic outcomes *in-cellulo* has long been employed in drug discovery as a way of measuring the effectiveness of medicinal chemistry optimizations. Additionally, proximal biomarkers have played increasingly important role in correlating the effect of target engagement on a small-molecule’s efficacy. However, in the absence of proximal reporters, distal biomarkers have the potential of confounding the correlation between target engagement and therapeutic efficacy and cannot tease apart the possibility of off-target interactions and the resultant toxicity (Davies et al., manuscript under review). Thus, the need for assessing direct and selective target engagement *in-cellulo* is highly desirable to provide insights into the extent of target protein’s inhibition required for the desirable phenotypic outcome. Further, if the method has the resolution to monitor a targeted set of homologous proteins, it becomes a powerful tool in the hand of a medicinal chemist. This feature confers an additional advantage of understanding cell-types based differences in target engagement during early drug-discovery (i.e. uptake, metabolism, distribution and so forth). Additionally, the cellular target-engagement assay has to be reasonably high-throughput for it to support early drug discovery.

**Figure 5.**
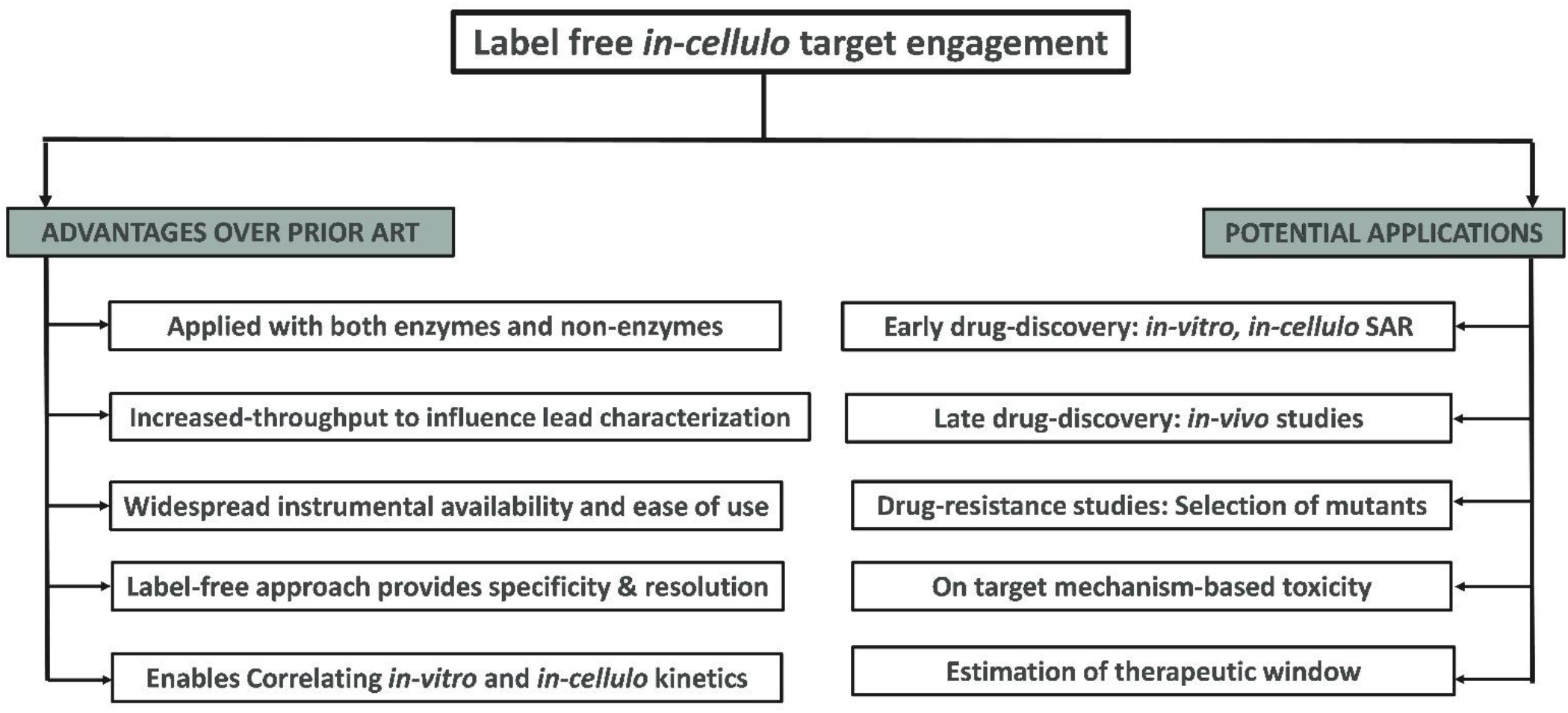
Flowchart showing the advantages of this method over similar approaches and potential application in early drug discovery.

The method espoused in the article has potential advantages over other orthogonal methods that have been employed to understand *in-cellulo* target engagement, i.e. FRET-based labelled assays (NanoBRET™)^27–29^ and/or CETSA^30–31^. NanoBRET™ systems necessitate the engineering of appropriate donor/acceptor pairs while CETSA cannot be applied with proteins that result in complex or, occasionally, no melting curves. On the contrary, label-free approaches that rely on chemoproteomic strategies, such as Activity Based Protein Profiling (ABPP)^32^, provide a very reliable estimate of *in-cellulo* target engagement translating into therapeutic outcomes. Although ABPP offers great resolution and sensitivity, especially for low abundance targets, they lack throughput or require high-end analytical infrastructure. The current study, by employing UPLC-MRM MS platform, has shown high sensitivity and increased throughput with a run time of 7.3 min per sample. Compared to nanoLC, this has 4-12 fold enhancement in throughput and will enable the interrogation of more samples (across various cell lines, dose & time dependent experiments) within a given time-frame, supporting iterative SAR across cell lines and timepoints. Towards the lower limit, this throughput translates into screening of 25 compounds in 30 hours as against 200 hours for nanoLC based approaches. This throughput will be reflected as quicker decision making during method development and quicker timeframe of iteration during SAR optimizations. Additionally, this will ensure that the hit validation approaches would be statistically robust, facilitating increased depth of mechanistic characterization. Another major advantage of the UPLC-QQQ based workflow is the lower premium on maintenance, availability and training compared to high-resolution instruments. However, low target abundance and/or limited sample availability can hamper the use of this platform for characterizing target-engagement. Having said that, a major limitation of all targeted proteomics approaches towards quantification of target engagement for irreversible has to do with the availability of unique proteotypic peptides and their ability to ionize in MS for reliable detection.

As alluded to earlier, a major advantage of the method is its label-free nature. Conventionally, kinetic parameters are estimated for enzymes that are amenable to assay design with a suitable means of measuring the substrate to product conversion. However, with increasing complexity of the drug discovery landscape, a lot of emerging targets are non-enzymes. This study assess the intracellular target engagement of one such non-enzymatic target by estimating parameters like *k_mod_*/*K_I_* (as a proxy for *k_inact_*/*K_I_*) making this method applicable to a wider range of proteins (**Figure 5**). This is a substantial advancement from the perspective of early drug discovery. Customarily, in drug-discovery, parameters like IC_50_ have been used to establish a correlation with *in-vitro* potency estimates. However, IC_50_ is a substrate concentration and time-dependent parameter which becomes concerning for a non-equilibrium modality of inhibition. Intracellular estimation of IC_50_ values are further constrained by non-Michaelian aspects such as liquid-liquid phase separation constraining diffusion and local sequestration, inhomogeneity, parameter not constrained by initial velocity, and so forth. Estimation of the second order kinetic parameter such as *k_mod_*/*K_I_* addresses this to some extent. Though the parameter is substrate concentration dependent, it is time-independent and yields an approximation of the inhibitor potency intracellularly under steady-state substrate concentration.

In a lot of cases, as exemplified in this study with KRAS^G12C^, specific targeting of the disease relevant protein and ability to selectively discriminate between the mutant and the wild-type is an essential prerequisite for attaining the right therapeutic window (**Figure 5**). In this case, the probe would have to specifically target the mutant variant (KRAS^G12C^) without affecting the WT KRAS, other variants of KRAS (G12D, G13C and G13D) and other RAS (HRAS, NRAS) proteins. We have demonstrated convincingly in this study that the small-molecule is highly specific to the variant of interest in this cell-line and does not interfere with the wild-type and homologs. Though the method was applied here as a means of assisting early drug discovery, we envision that this assay would be equally useful when the covalent molecules advance through the pipeline to *in-vivo* models/xerographs to tease apart the different levels of target engagement in different organs together with the off-target profiling, including for safety studies. An additional field of study would be to establish a link between target engagement and development of drug tolerance as a result of Darwinian selection of mutant variants that are drug resistant (**Figure 5**).

Thus, the current workflow demonstrates a way of proceeding from hit generation to lead-optimization that fully integrates the aspect of *in-cellulo* target engagement and kinetic parameter estimation. Employing various bio-analytical tools (LC and MS), this workflow strives to quantify the endogenous target engagement, including validating the conjugation of covalent compound with intact protein, helping choose the appropriate peptides from recombinant protein for analysis, confirming peptide sequence, endogenous sample preparation, optimizing LC and MS parameters and retrieving *in-cellulo* kinetic parameters. We envision that this workflow can be applied to different targets amenable to the covalent modality of inhibition and can be placed right after the biochemical hit identification and before *in vivo* characterization. This will help guide medicinal chemistry efforts in translating potent *in-vitro* covalent probes to efficacious *in vivo* equity and decreasing the turnaround time of SAR. Additionally, the technical advance presented in this study will enable the wider implementation of this workflow spanning labs across academia and industry employing the commonly available and less expensive QQQ systems.

### Conclusion and future perspective

This manuscript, to the best of our knowledge and belief, is the first ever demonstration of a high-throughput quantitative methodology that is capable of assessing endogenous on-target engagement and assessing *in-cellulo* kinetics for covalent probes in early drug discovery. While the methodology was implemented for the oncogenic target KRAS^G12C^, we posit that it is applicable to a broad range of targets encompassing enzymes, structural proteins and signalling molecules that have a nucleophilic cysteine modulating their activity and interactome.

**Table 2.**
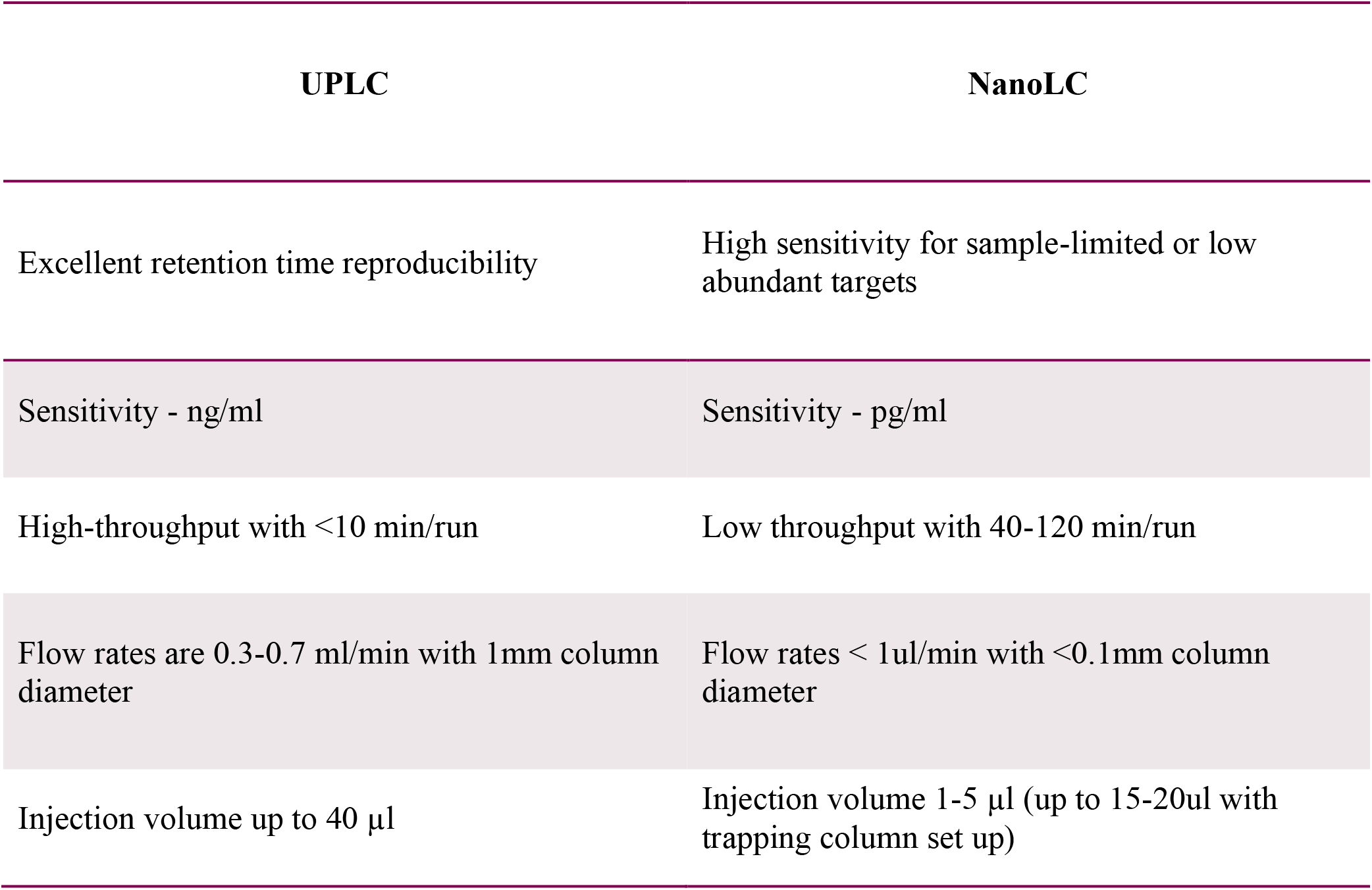
Advantages and disadvantages of nanoLC versus UPLC.

| UPLC | NanoLC |
| --- | --- |
| Excellent retention time reproducibility | High sensitivity for sample-limited or low abundant targets |
| Sensitivity - ng/ml | Sensitivity - pg/ml |
| High-throughput with <10 min/run | Low throughput with 40-120 min/run |
| Flow rates are 0.3-0.7 ml/min with 1mm column diameter | Flow rates < 1ul/min with <0.1mm column diameter |
| Injection volume up to 40 µl | Injection volume 1-5 µl (up to 15-20ul with trapping column set up) |

## Abbreviation

RF-MS: RapidFire Mass spectrometer
UPLC: Ultra performance liquid chromatography
MRM: Multiple reaction monitoring
LC: Liquid chromatography
DMSO: Dimethyl sulfoxide
WT: Wild-type
QQQ: Triple quadrupole
Q-TOF: Quadrupole time-of-flight
DTT: Dithiothreitol
RP: Reverse Phase
pM: PicoMolar
UHP: Ultra High Purity

## Acknowledgements

We would like to thank Maria Flocco, Liz Roberts and Chris Philips for their support and encouragement to carry out this work. Further, we would like to acknowledge the discussions that we had with Mohammad Pirmoradian, Ganesh Kadamur and Xiang Zhai that has improved the manuscript substantially. Protein Sciences Department, Compound Management and Mass Spectrometry facility are acknowledged for providing critical resources to carry out this study.

## Supplemental Materials

**Figure S1.**
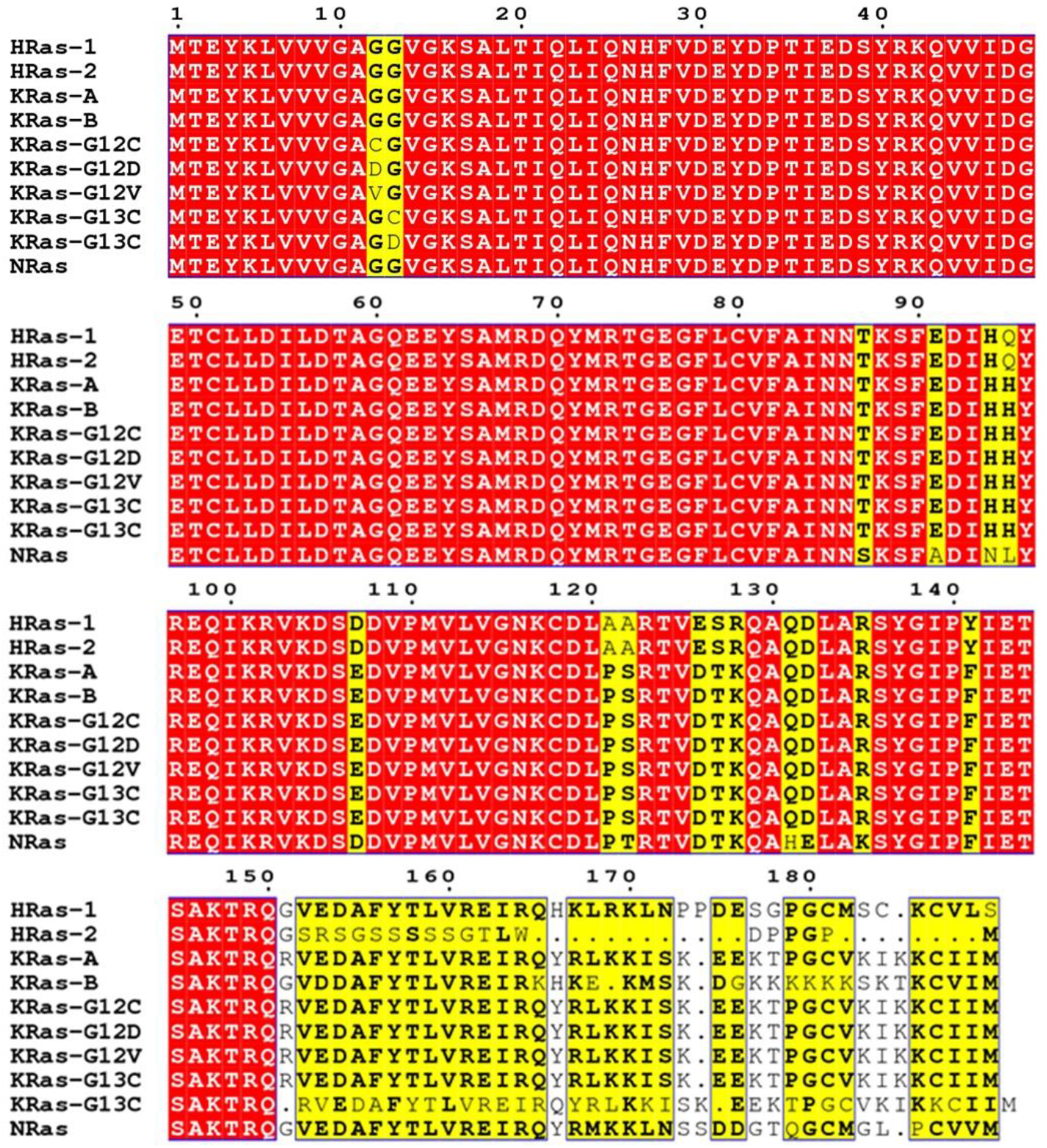
Connected to main **Figure 2**. Multiple sequence alignment of different RAS isoforms showing high degree of residue conservation in the N-terminus of the protein within the family. Codon 12 and codon 13 are indicated by yellow columns showing poor conservation. The sequence accession numbers are NRAS, NP_002515.1; KRAS-A, NP_001356715.1; KRAS-B, NP_001356716.1; HRAS-1, NP_001123914.1; HRAS-2, NP_789765.1; The MSA was generated with T-Coffee and the image was generated using ESPript 3.

**Figure S2.**
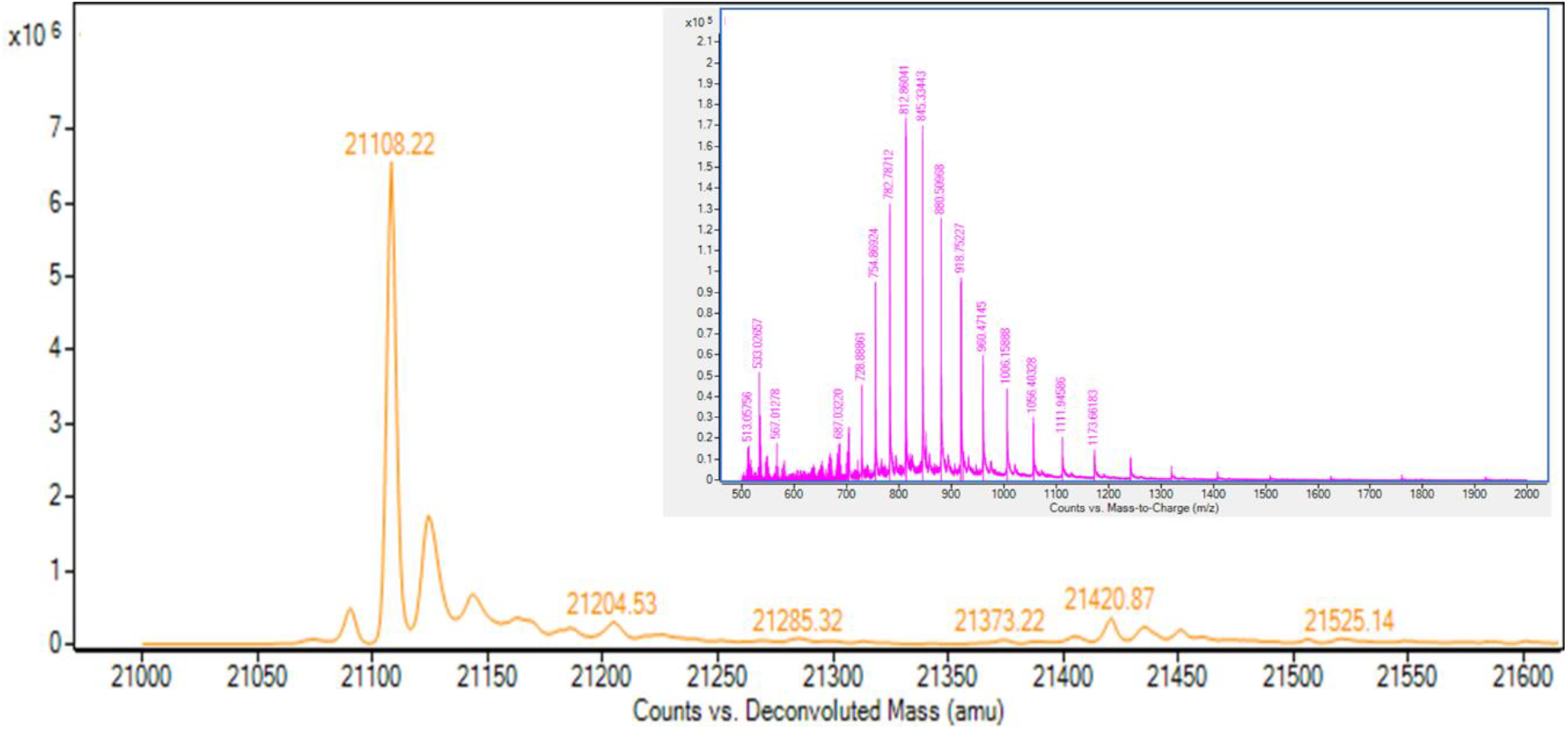
Connected to main **Figure 2C.** Deconvoluted intact mass and multiple charged raw MS spectrum of the KRAS^G12C^ intact protein injected at 1μM concentration.

**Figure S3.**
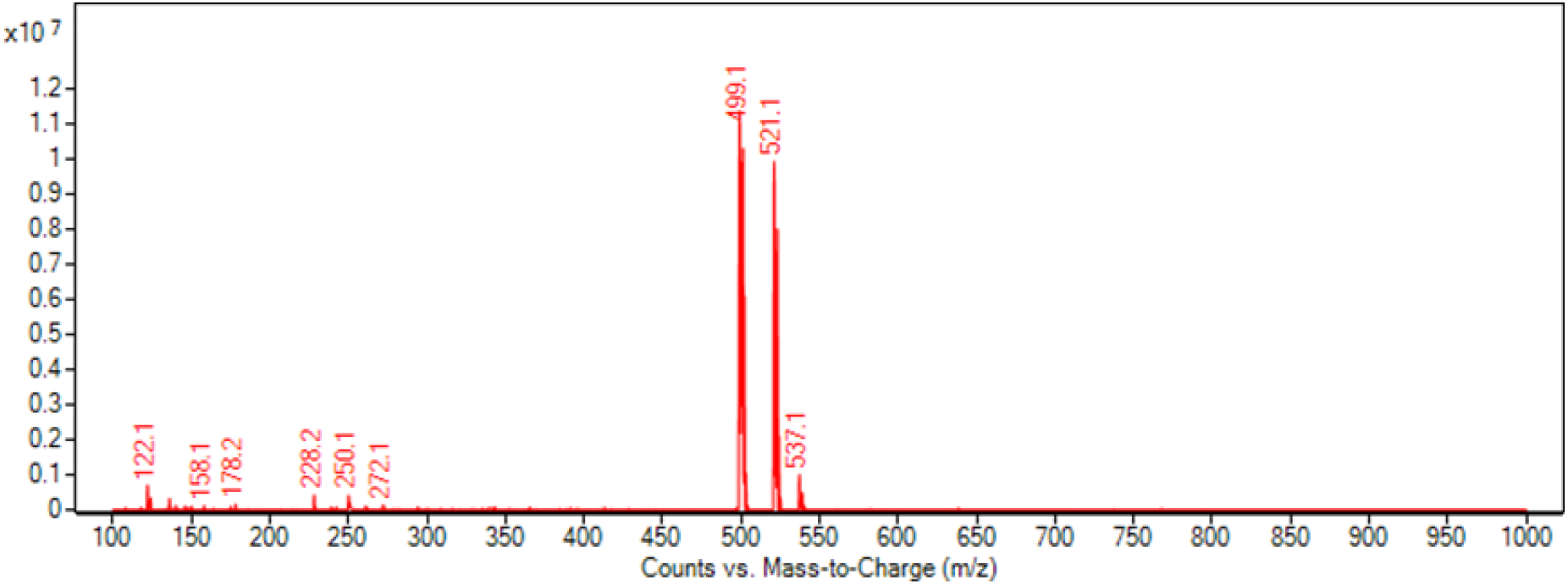
Connected to main **Figure 2C.** Mass spectrum confirming the mass of the covalent compound, compound 25 (*m/z* 499.1, M+H) along with the sodium adduct (*m/z* 521.1 M+Na) and potassium adduct (*m/z* 537.1 M+K)

**Figure S4.**
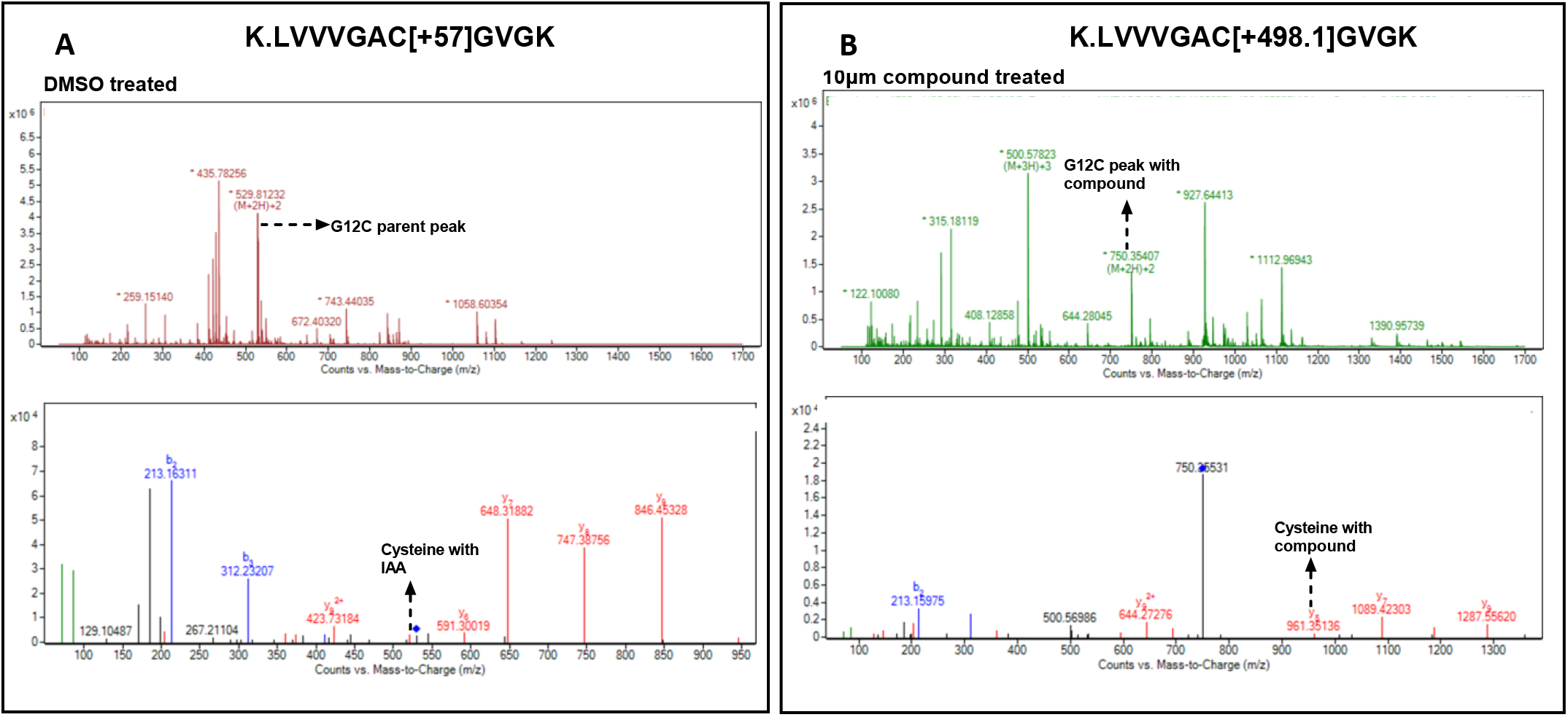
Connected to main **Figure 2D.** Peptide mapping of KRAS^G12C^ peptide. G12C peptide fragment spectra for the **A.** DMSO treated (modified with Iodoacetamide, IAA) and **B.** the corresponding compound treated peptide, validating the conjugation of the compound with the target cysteine. Both N-terminal b-ions and C-terminal y-ions are shown. *m/z* 1058.6 (*m/z* 529.8 doubly charged) corresponds to IAA modified and *m/z* 1499.7 (*m/z* 750.35 doubly charged) corresponds to compound 25 modified G12C peptide.

**Figure S5.**
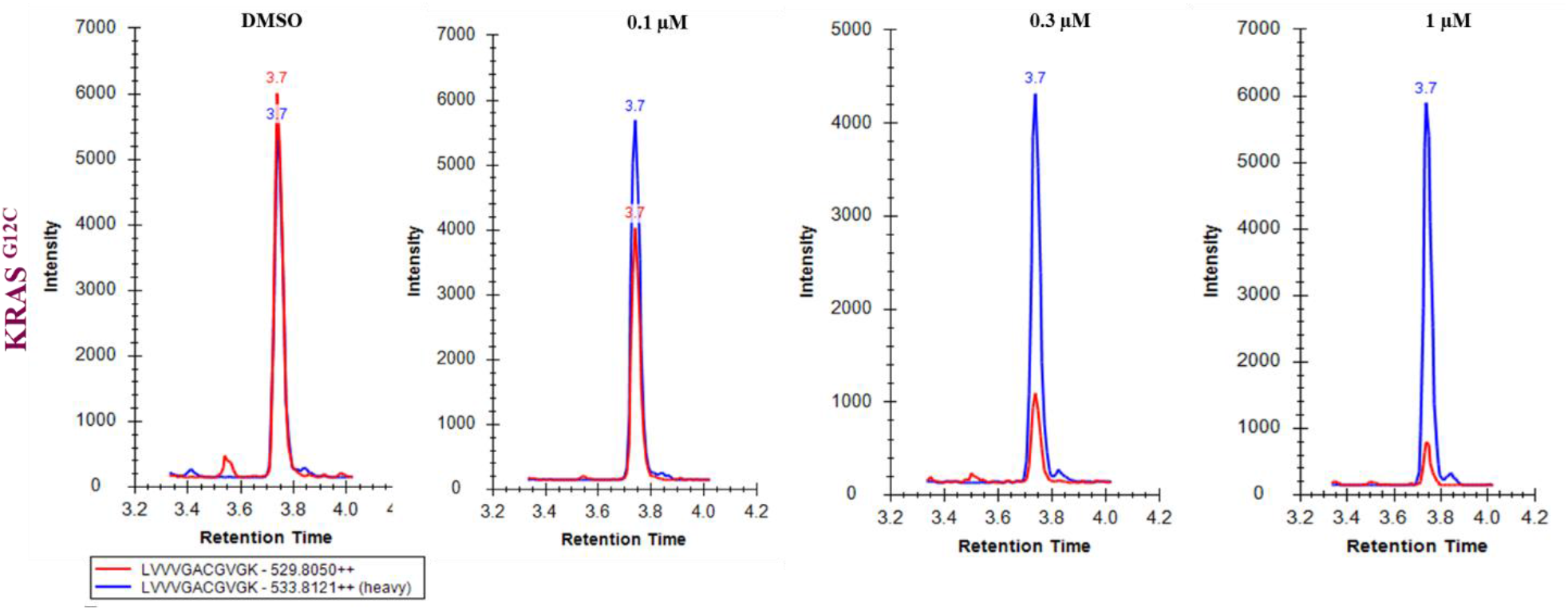
Connected to **Figure 4B**. Chromatograms displaying the comparison of the dose dependence of endogenous light protein (red) versus the spiked heavy standard (blue).

**Figure S6.**
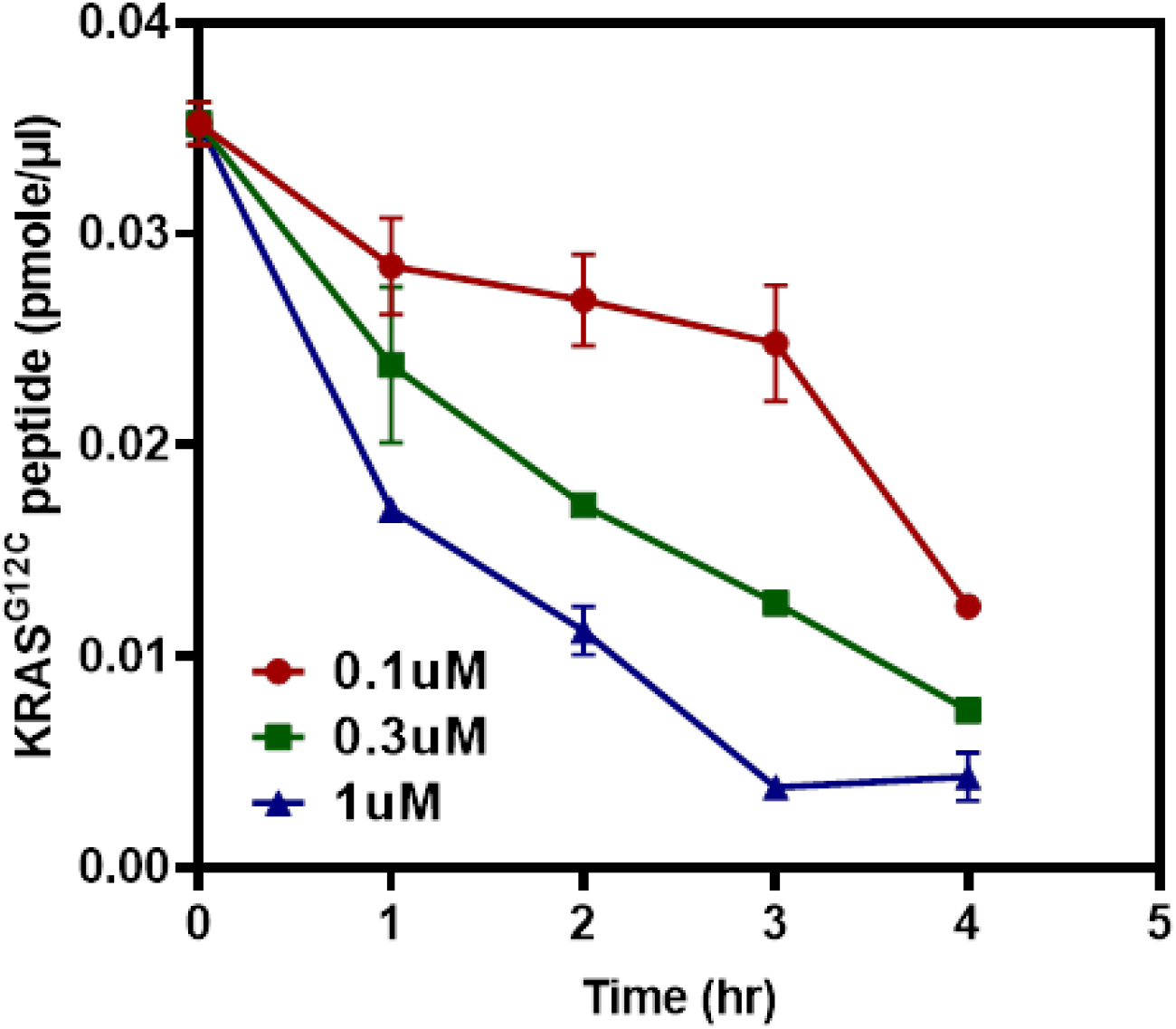
Connected to **Figure 4C**. Reduction in KRAS^G12C^ peptide as a function of increasing dose of compound 25 and across various time points.

